## Supplemental Table 4 for "Mechanism of life-long maintenance of neuron identity despite molecular fluctuations"

| **Supplementary Table.** ssODNs and guides used in this research. | | |
| --- | --- | --- |
| **ssODN** | **Allele** | **Sequence** |
| 3414 | *che-1::GFP::AID* | TGCTATACGAAGTTATTTTAAACTTACCTTTCCCTTCACGAACGCCGCCGCCTCCGGGCCACCGCTTGATTTTTGGCAGGAAACCATCACGTTCTTCCGGTATGATCGCACCGGTGGCCATCCCACAACTTGTGCCTTGGCCGGAGGTTTGGCTGGATCTTTAGGCATGGATTGAAAGTACAGATTCTCCTTGTAGAGCT |
| 3416 | *(ASEgcy-22 +flanks)p::che-1* | TGTTCTCTAAAATTGAAAAATAAGGAATCAGAATGTTTTTATATATTTTCAGTTATATCACATTTTACGAGCATTTATGTGCCCCACTTCGAATTTGAAGGGCTTCCCATAGACGCGTGATTTTATTTCGAATTTCGAGATTTCGAGAAAATTAAACCGTTTTTTTTCTCTTCTTCTTCAGCTCATCATCGAATCTCACC |
| 3417 | *(ASEgcy-22)p::che-1* | AAGGAATCAGAATGTGTAGGAAGTTGTTAGTAAGTGAAGCCCTTCAATTCATAGATAAGAAGAAACTACGGTATTCGGATTA |
| 3420 | *(deltaHD)p::che-1* | AGCTGAAGAAGAAGAGAAATTGGGAGAGAAGAGATGAATACCGTAGTTTCTTCTTATCTATGAAAATTGTGGCT |
| 3415 | *(ASEche-1+flanks)p::gcy-22* | ATTTTCCTGGTTATCTGACGTTAATTCAGTGATTCAATTGTTCTCTAAAATTGAAAAATAAGGAATCAGAATGTGTAGGAAGTTGTTAGTAAGTGAAGCCACAATTTTCATAGATAAGAAGAAACTACGGTATTCGGATTAAAATCTCTTCTCTCCCAATTTCTCCCTACGTCTTTGAATATTATAGGATTTCACAAAAT |
| 3418 | *(ASEche-1)p::gcy-22* | ATCTCGAAATTCGAAATAAAATCACGCGTCTATGGGAAGCCACAATTATTCGAAGTGGGGCACATAAATGCTCGTAAAATGT |
| 3419 | *(ASEche-1+flanksgcy-22)p::che-1* | TGTTCTCTAAAATTGAAAAATAAGGAATCAGAATGTTTTTATATATTTTCAGTTATATCACATTTTACGAGCATTTATGTGCCCCACTTCGAATAATTGTGGCTTCCCATAGACGCGTGATTTTATTTCGAATTTCGAGATTTCGAGAAAATTAAACCGTTTTTTTTCTCTTCTTCTTCAGCTCATCATCGAATCTCACC |
| **guide** | **Allele** | **Sequence** |
| g2 | *che-1::GFP::AID* | TCTGTACTTTCAATCCGGAA |
| g51 | *(ASEgcy-22 +flanks)p::che-1*  *(ASEche-1+flanksgcy-22)p::che-1* | GCTGAAGAAGAAGAGAAATT |
| g52 | *(ASEgcy-22 +flanks)p::che-1*  *(ASEche-1+flanksgcy-22)p::che-1* | AATAAGGAATCAGAATGTGT |
| g53 | *(ASEgcy-22)p::che-1* | CTTCTTATCTATGAAAATTG |
| g54 | *(deltaHD)p::che-1* | TAAGAAGAAACTACGGTATT |
| g55 | *(ASEche-1+flanks)p::gcy-22* | ATATTCAAAGACGTAGGTGT |
| g56 | *(ASEche-1+flanks)p::gcy-22* | CGTTAATTCAGTGATTCATA |
| g57 | *(ASEche-1)p::gcy-22* | TGCCCCACTTCGAATTTGAA |
